# Co-transcriptional translation amplifies mRNA noise in *Escherichia coli*

**DOI:** 10.1101/2023.10.25.563316

**Authors:** Sora Yang, Soojin Park, Jung Bae Son, Seunghyeon Kim, Soojung Yi, Gayun Bu, Nam Ki Lee

## Abstract

The variability in mRNA expression among isogenic cells exposed to identical environments is inherent. This variability originates from the inherent stochasticity of all processes underlying transcription. Although transcription and translation can occur simultaneously on the same mRNA molecule in bacteria, it is not well understood whether and how co-transcriptional translation contributes to variability in mRNA expression. Here, we studied the contribution of co-transcriptional translation to mRNA noise in *E. coli* cells. Using a transcription system physically decoupled from translation, we investigated the effect of ribosome binding to mRNA transcripts on variability in mRNA expression. We found that the propagation of RNAP noise to the mRNA level was increased by ribosome binding, leading to larger variations in the mRNA levels. We further demonstrated that ribosome binding increased the transcription initiation rate, resulting in the promoter becoming susceptible to RNAP noise. Co-transcriptional translation amplified transcriptional noise and modulated transcriptional bursting kinetics in bacterial cells.

## Introduction

Variability (noise) in mRNA expression among isogenic cells exposed to the same environment is inevitable. This phenomenon limits the accuracy of gene regulation but plays an important role in diverse cellular processes, such as cell fate decision, state switching, and bacterial survival (Urban and Johnston, 2018). This variability arises from the inherent stochasticity underlying all transcription processes (intrinsic noise) and cell-to-cell variations in the cellular environment (extrinsic noise) (Elowitz et al., 2002). Most current studies have assumed that the transcription initiation process is a major source of mRNA noise and focused on how transcription initiation including promoter sequence (Larsson et al., 2019; Ozbudak et al., 2002; So et al., 2011), transcription factor binding (Gronlund et al., 2013; Hensel et al., 2013; Jones et al., 2014; Sepulveda et al., 2016; To and Maheshri, 2010), and initiation processes by RNAP (Park et al., 2018; Yang et al., 2014) or sigma factors (Engl et al., 2020) affects variability in gene expression. Recent studies have investigated the effects of other processes such as transcription elongation dynamics (Kim and Jacobs-Wagner, 2018), DNA replication (Peterson et al., 2015), and mRNA degradation (Baudrimont et al., 2019; Kim and Jacobs-Wagner, 2018) on gene expression noise.

In living bacterial cells, ribosomes bind to mRNA transcripts immediately after a ribosome binding site (RBS) is generated during RNAP elongation. Electron microscopy has revealed that translating ribosomes closely follow the transcribing RNAP (Miller et al., 1970). Recently, the structural interactions between ribosomes and RNAP have been extensively studied (Demo et al., 2017; Fan et al., 2017; Kohler et al., 2017; Wang et al., 2020; Webster et al., 2020). Consequently, translating ribosomes can affect transcriptional dynamics. A well-known example is that the transcription elongation rate is tightly controlled by the translation rate (Proshkin et al., 2010). Our single-molecule *in vivo* imaging study revealed that transcription is spatially coupled with translation and functional coupling in live *E. coli* cells (Yang et al., 2019). However, it is not clear whether and how translation contributes to variability in mRNA expression.

Here, we investigated the contribution of translation to mRNA noise. Using a T7 RNAP system that is physically decoupled from translation, we focused on the effect of ribosome binding to mRNA transcripts on mRNA noise. We simultaneously measured RNAP concentration variation and mRNA copy numbers from specific promoters in individual *E. coli* cells and quantitatively investigated the contribution of ribosome binding to mRNAs at the variability in mRNA level. We demonstrated that the propagation of RNAP noise to mRNA noise was increased by ribosome-binding events, leading to larger variations in mRNA levels. These results are reconciled by a mechanism in which co-transcriptional translation increases the transcription initiation rate, and the probability that other free RNAPs bind to the promoter increases; therefore, the promoter becomes susceptible to RNAP noise. Our data suggest that translation, as a post-transcriptional process, influences pre-transcriptional processes in live bacterial cells.

## Results

### Simultaneous measurement of RNAP concentration and mRNAs in individual *E. coli*

Transcription and translation are intricately coordinated in bacterial cells. The transcriptional elongation rate is controlled by the translation rate (Proshkin et al., 2010). Moreover, blocking translation leads to premature transcriptional termination, and recent structural and biochemical studies have offered insights into the direct molecular interaction between RNAP and ribosomes *in vitro* (Stevenson-Jones et al., 2020; Wang et al., 2020; Webster et al., 2020). However, the mechanistic cooperativity between RNAP and ribosomes *in vivo* has remained elusive. To circumvent complications arising from the unknown cooperation between endogenous *E. coli* RNAP and ribosomes in cells, we used our recently developed T7 RNAP-transcription imaging system (Yang et al., 2019; Yang et al., 2014) (Fig. 1a). Given that T7 RNAP is a single-subunit enzyme, it lacks a known physical link with the endogenous *E. coli* ribosomes. Because the elongation rate of T7 RNAP is more than five times faster than that of ribosomes, T7 RNAP can be mechanically decoupled from ribosomes. However, translation can still occur in an mRNA when T7 RNAP produces mRNA transcripts. Therefore, we could focus on the effect of co-transcriptional translation using the T7 RNAP transcription system. Furthermore, using this system, the concentration of T7 RNAP can be controlled and measured in individual cells. We also monitored the regulated gene activity of the T7 promoter by measuring the downstream mRNA or protein expression levels. Analysis of single-cell data allowed us to identify the propagation of RNAP noise to downstream mRNA noise and the contribution of RNAP concentration to the stochastic activity of the promoter.

**Figure 1.**
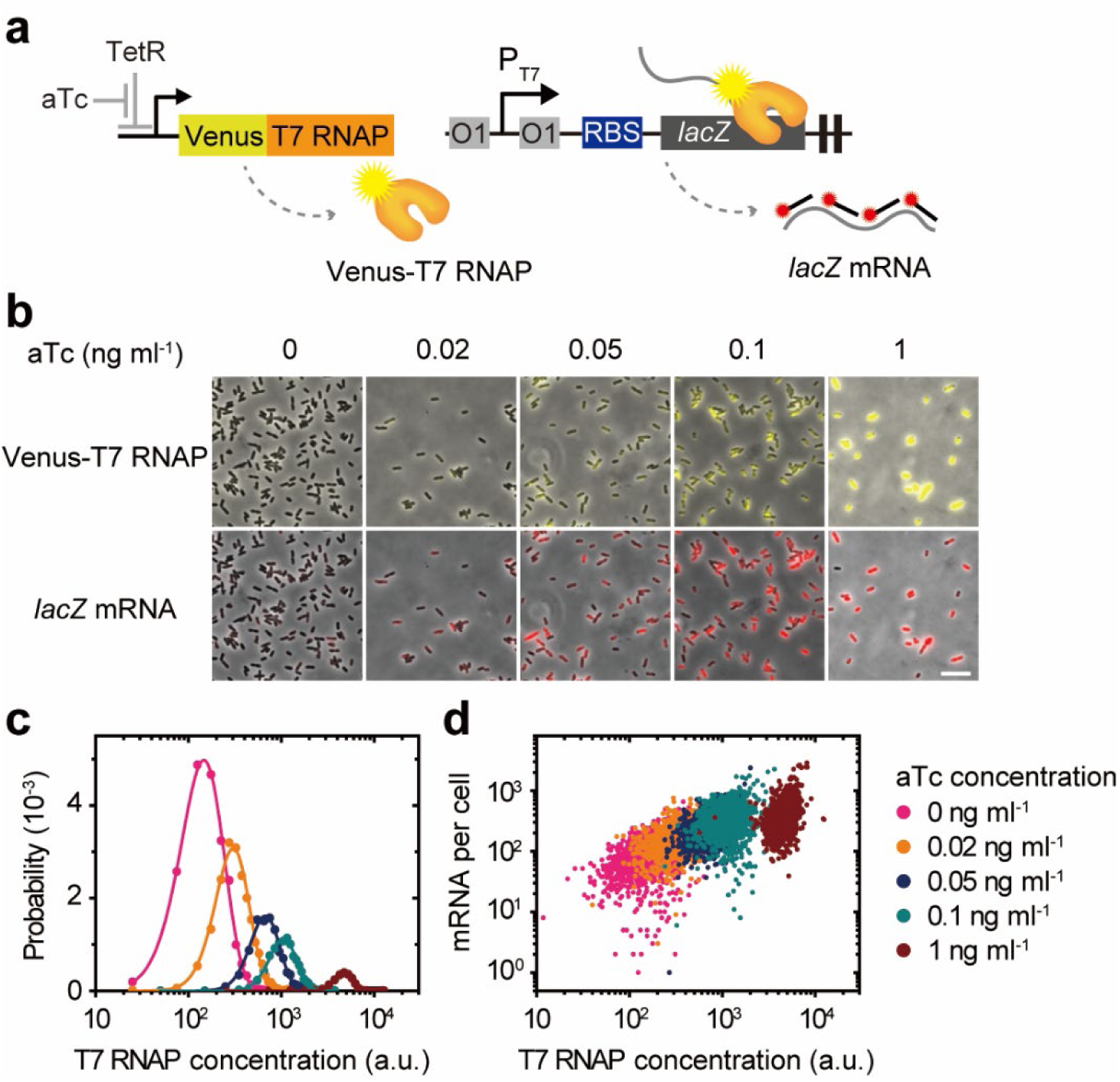
Simultaneous measurement of RNA polymerase concentration and mRNA copy numbers in individual *E. coli* cells. (a) T7 RNAP-driven expression of *lacZ* mRNA in individual cells. Venus-T7 RNAP is expressed from a tetracycline promoter in the plasmid, pNL001. Varying the concentration of inducer aTc controls the expression level of Venus-T7 RNAP. The *lacZ* gene is expressed under T7 promoter, which is replaced with the endogenous *lac* promoter. Two *lac* operators (O1) are inserted, one upstream and one downstream of the T7 promoter. To ensure full expression of *lacZ*, 1mM isopropyl β-D-1-thiogalactopyranoside (IPTG) was added. (b) Representative overlay images of the phase contrast and fluorescence images at each aTc concentration. Scale bar, 10 μm (c) Steady-state distributions of Venus-T7 RNAP concentration under different aTc concentrations. Solid lines are fits to gamma distributions. (d) The number of *lacZ* mRNAs in individual cells is plotted against T7 RNAP concentration under different aTc concentrations. Each dot represents individual cell.

We replaced the endogenous *lac* promoter with the T7 promoter. The plasmid expressed T7 RNAP tagged with the yellow fluorescent protein Venus (pNL001), which allowed us to control and measure the T7 RNAP concentration in cells (Fig. 1a). The *lacZ* mRNA driven by T7 RNAP was quantified using single-molecule fluorescence *in situ* hybridization (smFISH) (Skinner et al., 2013). The probe set for detecting *lacZ* mRNA comprised of 50 fluorescently-labelled oligonucleotides against *lacZ* transcripts. Venus-T7 RNAP and *lacZ* mRNA levels in individual cells were simultaneously measured using epifluorescence microscopy after hybridization with the probe set to fixed cells (Fig. 1b). The distributions of Venus-T7 RNAP concentration were obtained at each inducer, anhydrotetracycline (aTc), concentration and fitted to gamma distributions (Friedman et al., 2006) (Fig. 1c). The expression levels of *lacZ* mRNA were dependent on T7 RNAP concentration (Fig. 1d).

### RNAP noise propagation to the downstream gene expression

Our T7 RNAP transcription imaging system provides a platform for investigating RNAP noise propagation to downstream gene expression (Yang et al., 2014). Using the data depicting RNAP-concentration-dependent mRNA expression levels in individual cells (Fig. 1d), we manipulated RNAP noise while maintaining a fixed mean RNAP concentration, which allowed us to examine how RNAP noise variation propagates downstream. We generated cell distributions with a predetermined mean and standard deviation (s.d.) of RNAP concentration from the total cell population (Fig. 2a, left). We combined all cells from Fig. 1c (∼ 20,000 cells) and then randomly selected 300 cells from the total cell data, in which the sub-sampled cells followed a gamma distribution with a fixed mean and s.d. to mimic the natural expression system (Fig. 2b, middle). We confirmed that randomly selected cell populations showed a similar trend in T7 RNAP concentration-dependent mRNA expression levels under different aTc induction conditions (Supplementary Fig. 1). The mRNA distribution obtained from the selected subset of cells was well-described by a negative binomial distribution predicted by the two-state model (Fig. 2a, right) (Paulsson and Ehrenberg, 2000; So et al., 2011). Conducting this process over a range of T7 RNAP concentrations allowed us to establish the relationship between RNAP noise (η^2^) and mRNA noise (η^2^) at each constant T7 RNAP concentration (Fig. 2b); mRNA noise increases linearly with the increase in RNAP noise and the propagation efficiency of RNAP noise to mRNA noise (slope in Fig. 2b) is inversely proportional to the RNAP concentration. This behavior is consistent with the RNAP noise propagation aspect of protein noise in a previous study (Yang et al., 2014); however, the propagation efficiency of RNAP noise was higher at the mRNA level than at the protein level (Fig. 2c), showing that cell-to-cell variability in RNAP concentration propagates to each downstream level and, at the protein level, is attenuated by 40 % of the mRNA level (Fig. 2d). mRNA transcripts produced by RNAP undergo multiple rounds of translation, and their lifetimes are much shorter than those of proteins. This phenomenon may attenuate the propagation of variations in the RNAP concentration at the protein level.

**Figure 2.**
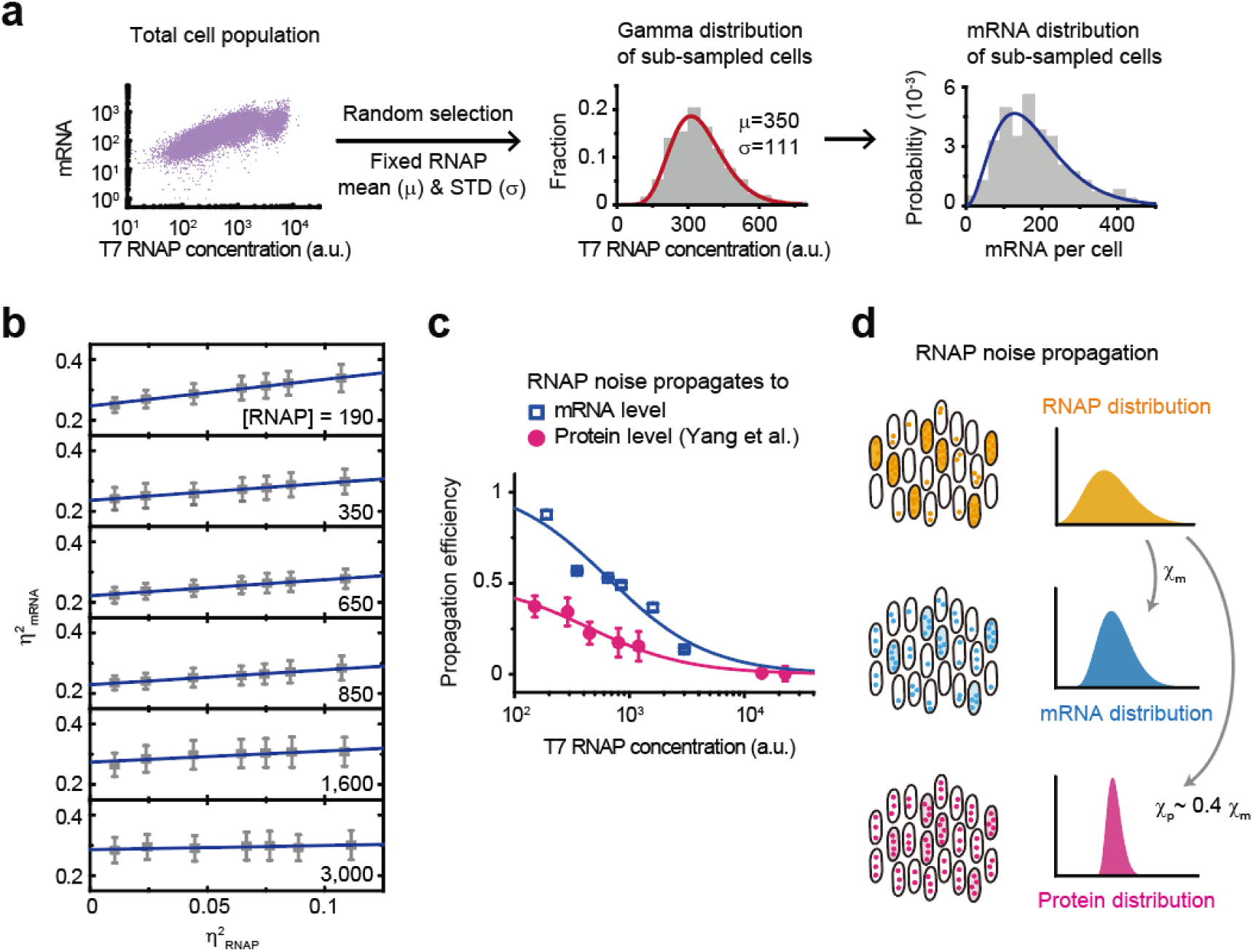
RNAP noise propagation to the downstream gene expression. (a) Random generation of cell distributions with a pre-determined mean and s.d. of the RNAP concentration from the total cell population. Left, total cell population obtained by combining all cells from Fig. 1c (∼ 20,000 cells). Each point represents single-cell data. Middle, distribution of the subset of cells (300 cells) randomly selected from the total cell population to satisfy a given gamma distribution (solid line) with a pre-determined mean (μ) and s.d. (σ). Right, mRNA distributions in the sub-sampled cells. Solid line is fit to a negative binomial distribution. (b) mRNA noise of the sub-sampled cells depending on RNAP noise. Inset number: mean T7 RNAP concentration of the population of sub-sampled cells. Lines: linear fits. (c) Comparison of RNAP noise propagation efficiency to mRNA noise (the slope of the mRNA noise depending on the T7 RNAP concentration from b) with to protein noise (adapted from Yang et al. 2014 Nat Comm). Each graph fits the function of the unoccupied fraction of the promoter by RNAP, y=c/(1+Kx), where K and x denote the RNAP affinity for the promoter and T7 RNAP concentration, respectively. (d) Model of RNAP noise propagation to the downstream gene expression noise. The propagation efficiency of RNAP noise (susceptibility to RNAP noise, χ) is attenuated ∼ 40% of mRNA noise in protein noise.

As we used the RNAP concentration measured at a given time, the analysis did not consider the history of fluctuations in the RNAP concentration. However, in a previous study, we confirmed that the contribution of fluctuations in RNAP concentration over time was not significant at the protein level (Yang et al., 2014). In addition, the lifetime of the target mRNA was a few minutes (Supplementary Fig. 2), whereas the protein lifetime was typically longer than that of the cell cycle in *E. coli*. As a result, the mRNA level reflected a more recent history of RNAP concentration (a few minutes) than the protein level, showing that sampling analysis using cells with RNAP concentration and mRNA levels at a given time was still a valid approach for our study.

### Ribosome binding to mRNA increases mean and variability in mRNA expression level

To directly assess whether co-transcriptional translation affects variability in mRNA expression level, we inhibited translation by mutating the RBS in the 5’ untranslated region of the *lacZ*. The mutant (ΔRBS) exhibited several hundred times lower LacZ activity (Yang et al., 2019), and we compared the mRNA copy number distributions with (WT) and without RBS (Δ RBS) (Fig. 3a). The mRNA distributions of both strains were well described by a negative binomial distribution, and the mean mRNA levels were dependent on the T7 RNAP concentrations (Fig. 3b). However, when ribosome binding to mRNA was inhibited (ΔRBS), there was a significant decrease in mRNA level at each inducer concentration. The absence of translating ribosomes on mRNA transcripts may induce rapid mRNA decay by exposing mRNAs to the degradation machinery, although T7 RNAP is physically decoupled from *E. coli* ribosomes (Deana and Belasco, 2005). To identify differences in mRNA decay between the two strains, we measured the *lacZ* mRNA lifetime using quantitative reverse transcription polymerase chain reaction (qRT-PCR). When *lacZ* expression under the control of the T7 promoter was fully induced by IPTG, the expression was shut off by rapid exchange of the medium with no IPTG medium, and the decay of the mRNA level was measured using qRT-PCR (Supplementary Fig. 2). The difference in mRNA lifetime was approximately three times lower than the difference in mean mRNA levels at high T7 RNAP concentrations (approximately 4.5 times, Supplementary Fig. 3). This implies that T7 RNAP transcription may be affected by co-transcriptional translation in *E. coli,* despite the lack of physical interaction with ribosomes. We note that the fold change of the mean, defined as the ratio of the mean mRNA number with and without RBS, shows RNAP concentration dependence; ribosome binding to the mRNA transcripts affects T7 RNAP transcription in an RNAP concentration-dependent manner (Supplementary Fig. 3). This suggests that co-transcriptional translation affects RNAP concentration-dependent processes, such as transcription initiation, rather than RNAP concentration-independent processes.

**Figure 3.**
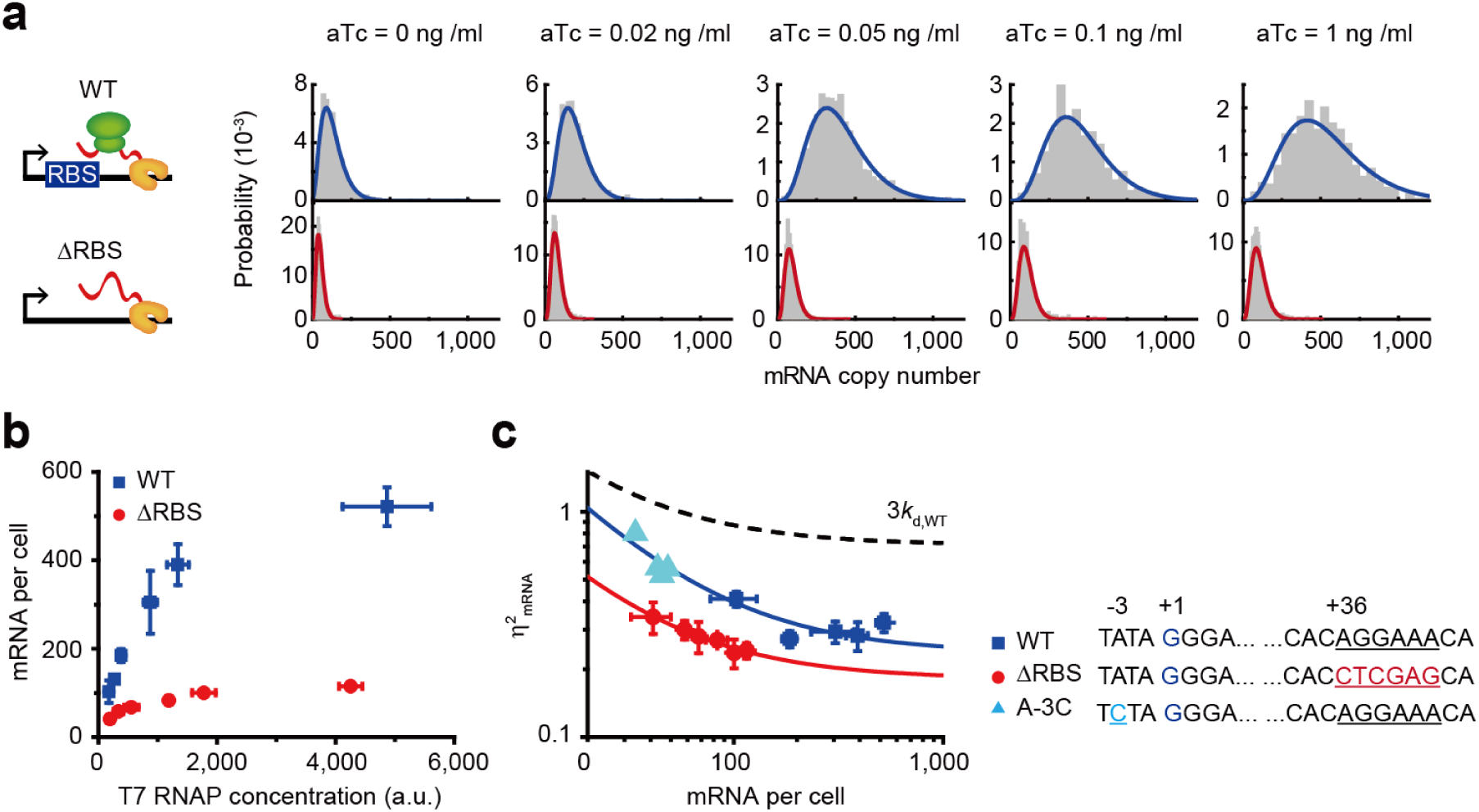
Ribosome binding to mRNA increases mean and variability in mRNA expression level. (a) mRNA distributions under different aTc concentrations with and without RBS. Solid lines are fits to negative binomial distributions. (b) T7 RNAP-driven mRNA expression levels under different aTc concentrations with and without RBS. (c) mRNA noise obtained from the mRNA distribution as a function of the mean mRNA. Solid curves fit y=a/x + b, where x = mRNA copy number; a = 16.05, b = 0.24 for WT; a = 6.67, b = 0.18 for ΔRBS. Error bars ± s.d., n = 3. Dotted line represents the cases of 3*k*_d_ for WT.

Next, we compared the mRNA noise with and without RBS (Fig. 3c). In both cases, mRNA noise decreased with an increasing mean mRNA copy number. However, for ΔRBS strain, there are some deviations from the basic trend of WT, indicating that ribosome binding to mRNA transcripts might be altering the gene expression behavior of WT. In other words, mRNA noise increases when translation is efficiently initiated through ribosome binding to mRNA transcripts. We noted that the increase in mRNA variability by co-transcriptional translation is dominated by altering stochastic activity of transcriptional regulation rather than of other processes such as mRNA degradation: (i) fast mRNA decay leads to increase in mRNA noise (Baudrimont et al., 2019), (Fig. 3c, dotted line) but ΔRBS shows lower mRNA noise. (ii) Even when mRNA expression levels were reduced by mutating the T7 promoter sequence (Imburgio et al., 2000), the mRNA noise remained higher than in the ΔRBS strain (Fig. 3c, light blue), suggesting that ΔRBS alters transcriptional activity in a different way than the T7 promoter mutant.

Although transcriptional regulation is influenced by the sequence of regulatory DNA, such as TATA box strength (Hornung et al., 2012) and the number of transcription factor binding sites (Raj et al., 2006; To and Maheshri, 2010), notably, the RBS is situated in a downstream region of the promoter. This region is not the regulatory DNA region for transcription initiation but rather the region through which the elongation-phase RNAP passes. Thus, although the mechanism of transcription regulation may not be identical to that of these cases, co-transcriptional translation clearly contributes to transcriptional regulation, even when physically uncoupled from translation.

### Ribosome binding to mRNA increases propagation of RNAP noise to mRNA noise

To better understand why ΔRBS exhibits lower mRNA noise than the WT, we conducted a similar analysis to assess the translation dependence of RNAP noise propagation efficiency. This was performed using data from T7 RNAP concentration-dependent mRNA levels obtained from ΔRBS cells (Supplementary Fig. 4), mirroring the approach shown in Fig. 2. For ΔRBS, the mRNA noise also increased linearly with increase in RNAP noise (Fig. 4a), but the propagation efficiency of RNAP noise to mRNA level was lower than that in WT (Fig. 4b). In other words, RNAP noise propagation efficiency increased with translation, leading to an increase in mRNA noise at the same RNAP concentration distribution (Fig. 4c). When the contribution of RNAP noise was removed (η^2^_RNAP_=0), mRNA noise converged to the same plateau in both strains. This suggests that the observed difference originated entirely from the difference in RNAP noise propagation between the two strains (Fig. 4c). The observed plateau may signify the existence of other extrinsic sources independent of RNAP concentration variations in our system (Taniguchi et al., 2010).

**Figure 4.**
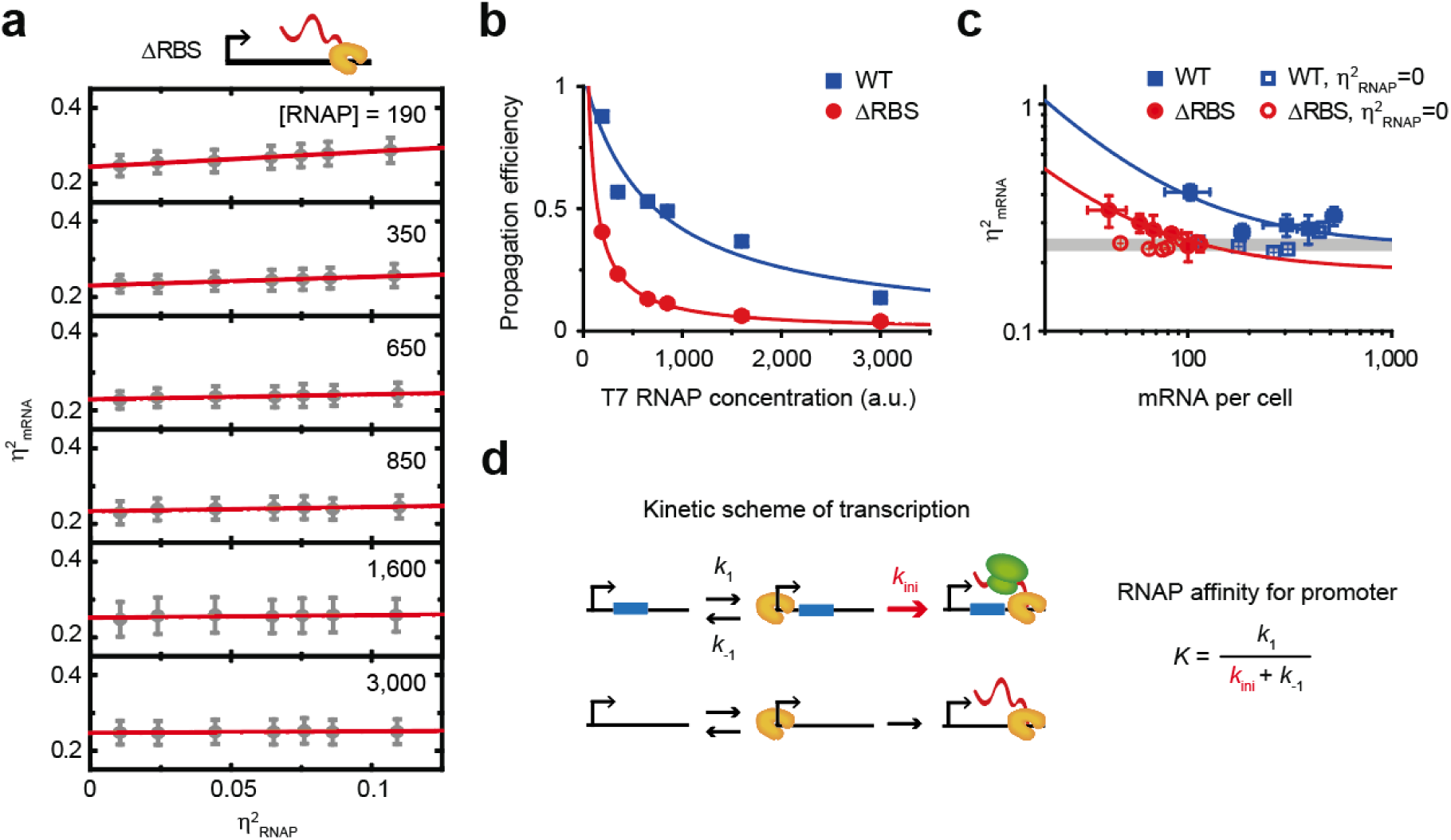
Ribosome binding to mRNA increases propagation of RNAP noise to mRNA noise. (a) mRNA noise of the sub-sampled cells depending on RNAP noise in the absence of RBS. Inset number: mean T7 RNAP concentration of the population of sub-sampled cells. Lines: linear fits. (b) The slope of the mRNA noise depending on the T7 RNAP concentration from (a). The data of WT from Fig. 2c (blue) were plotted for comparison. Each graph fits the function of the unoccupied fraction of the promoter by RNAP, y=c/(1+*K*x), where *K* and x denote the RNAP affinity for the promoter and T7 RNAP concentration, respectively. (c) Contribution of RNAP noise to mRNA noise. The mRNA noises at an η^2^_RNAP_=0 (open symbols) were obtained from the y axis intercepts resulting from linear fitting in a (ΔRBS, red) and Fig. 2b (WT, blue). For comparison, the mRNA noises shown in Fig. 3c were plotted with those at an η^2^_RNAP_=0. (d) Kinetic scheme of transcription. Translation increases transcription initiation (*k*_ini_), leading to lower RNAP affinity for promoter (*K*).

The dependence of RNAP noise propagation efficiency on RNAP concentration has been described as the fraction of promoters that are unoccupied by RNAP (Yang et al., 2014). Based on Michaelis-Menten enzyme kinetics, the steady-state rate of the transcriptional reaction represented in Fig. 4d is proportional to the fraction of the promoter occupied by RNAP, *θ* = *KN*_RNAP_ / (1 + *KN*_RNAP_), where *K* is defined by *K* = *k*_1_ / (*k*_ini_ + *k*_-1_) and *N*_RNAP_ is the RNAP concentration. For both strains, the propagation efficiencies well fit the function of the unoccupied fraction, 1 − *θ =* a / (1 + *KN*_RNAP_) (Fig. 4b) and the fitting allows extraction of *K*, described as the RNAP affinity for promoter. When translation was inhibited (ΔRBS), the promoter affinity for RNAP, *K*, increased about 10 times compared to WT. Because ΔRBS strain showed lower transcriptional activity (Fig. 3b), the increased affinity of RNAP to promoter can be explained by a decrease in transcriptional initiation rate, *k*_ini_. As the transcriptional initiation rate increases, the probability of other free RNAP binding to the promoter increases; thus, the promoter becomes more susceptible to RNAP noise.

To gain further support and confirm the decrease in the transcriptional initiation rate in the RBS, we quantified *in vivo* transcriptional kinetics using single-molecule imaging of YFP-T7 RNAP in living cells (Yang et al., 2019). By measuring the intensity of the fluorescent foci created when YFP-T7 RNAP transcribes the downstream gene of the T7 promoter as a function of time after transcription induction by adding the inducer IPTG, the transcriptional on-rate and elongation rate of T7 RNAP were obtained (Supplementary Fig. 5). As expected, the elongation rate of the T7 RNAP remained unaffected by translation. However, the transcriptional on-rate was slower by ∼ 33 % when translation was inhibited in ΔRBS. This aligns with the results of noise analysis, indicating that co-transcriptional translation enhances the transcriptional initiation process.

### Ribosome binding to mRNA regulates transcriptional burst size

In a two-state model of gene expression (Paulsson, 2005; Peccoud and Ycart, 1995; Shahrezaei and Swain, 2008), a gene switches between ‘on’ and ‘off’ states, and transcription can occur only during the ‘on’ state. The number of mRNA produced in the on-state (burst size) and switching rate between the two states (burst frequency) were defined in the model. By applying this stochastic model of gene expression, we determined the mRNA burst statistics (Fig. 5). In both strains, burst size exhibited a clear dependence on the RNAP concentration, whereas burst frequencies remained independent. These behaviors are consistent with the expectations based on the two-state model: during the ‘on’ state, successive binding of RNAP at high RNAP concentration led to the production of a large number of mRNAs, but had no major effect on the switching to the ‘on’ state of DNA (Fig. 5a and b). Notably, WT exhibited a higher burst size compared to ΔRBS (Fig. 5a), while the difference in the burst frequency did not show significant difference (Fig. 5b). These behaviors of the kinetic parameters can be explained by the above observation: the increased transcriptional initiation rate by translation should result in a higher extent of bursting, as reflected by a higher burst size.

**Figure 5.**
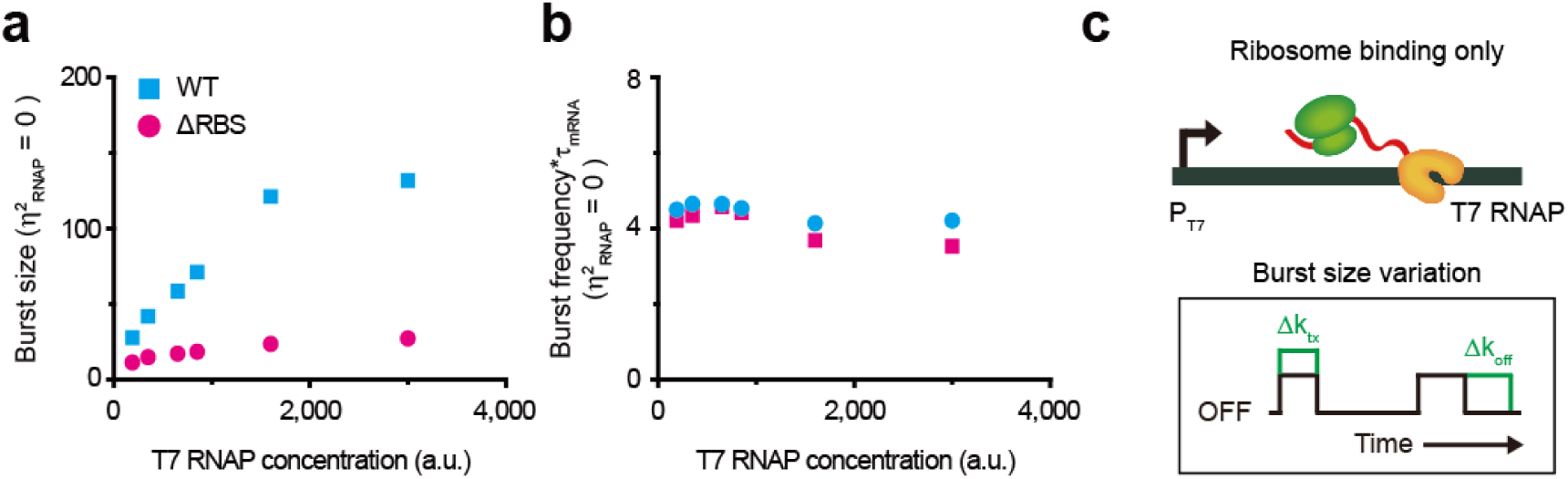
Ribosome binding to mRNA regulates transcriptional burst size. (a, b) We compared the burst kinetics, (a) burst size and (b) burst frequency, with those at an η^2^_RNAP_=0. Each quantity at an η^2^_RNAP_=0 were calculated from the y axis intercepts resulting from linear fitting in Fig. 2b and 4a. (c) Model of ribosome binding to mRNA contribution to transcriptional bursting kinetics.

## Discussion

Through single-cell measurements accompanied by quantitative analysis, we showed that the propagation of RNAP noise to the downstream mRNA level was enhanced when translation occurred on mRNA transcripts, leading to larger mRNA variability. Co-transcriptional translation induces an increase in the transcription initiation rate, which makes the promoter more sensitive to variations in RNAP concentration. Furthermore, we demonstrated that the propagation of RNAP concentration variation was attenuated by 40 % at the protein level compared to that at the mRNA level. Therefore, our findings are expected to be helpful for predicting noise propagation in gene networks and/or for designing noise-tolerant gene circuits.

Currently, there is no mechanistic understanding of how translation affects transcriptional initiation. Our finding that translation, even though it is physically uncoupled from transcription, affects transcriptional regulation suggests that the observed effect of translation involves a change in DNA topological features. Specifically, we previously showed that gene loci dynamically relocate during transcription, and the effect is potentiated by translation (Yang et al., 2019), suggesting that the dynamic relocation of gene loci may induce topological differences in the DNA promoter region, thereby affecting transcriptional regulation.

Notably, co-transcriptional translation may also be a possible source of the transcriptional bursting mechanism in bacteria. Transcription bursting in *E. coli* has been directly observed using the MS2 technique (Golding et al., 2005). DNA supercoiling has been experimentally proven as the mechanism that gives rise to cause bacterial transcriptional bursting (Chong et al., 2014). Our findings would provide new insights into the mechanisms underlying transcriptional bursting in bacteria.

## Author contributions

N.K.L. supervised the project; N.K.L and S.Y. designed the experiments; N.K.L., S.Y., and S.P. wrote the manuscript; S.Y. and S.P. performed the experiments and analyzed the data; and J.B.S, S.K, S.Y., and G.B. helped perform the experiments.

## Acknowledgments

This work was supported by the Creative-Pioneering Researchers Program of Seoul National University, and NRF-2023R1A2C2006606 and NRF-2020R1A5A1019141 of the National Research Foundation of Korea.

