## Supplementary materials for "Co-transcriptional translation amplifies mRNA noise in *Escherichia coli*"

### Supplementary information for
### “Co-transcriptional translation amplifies mRNA noise in *Escherichia coli*”

Sora Yang[1†], Soojin Park[1†], Jung Bae Son[1], Seunghyeon Kim[1,3], Soojung Yi[1], Gayun Bu[2], and Nam Ki Lee[1]

[1] Department of Chemistry, Seoul National University, Seoul 08826, Korea.

[2] Department of Physics, Pohang University of Science and Technology (POSTECH), Pohang 37673, Korea.

[3] Present address: Department of Physics, University of Illinois at Urbana–Champaign, Urbana, IL 61801, USA

† These authors contributed equally to this work.

#### Supplementary Materials and Methods

##### Bacterial strains

All bacterial strains and plasmids used are listed in Table S1.

##### Growth media and conditions

All strains from single colonies were grown overnight in 3 mL of LB at 37 ℃ with shaking. The overnight cultures were diluted 1:200 into M9 medium (supplemented with 0.4 % glucose, vitamins, and amino acids) in the presence of appropriated antibiotics (50 μg/mL Carbenicilin and 35 μg/mL Chloramphenicol for T7p_3.3kb, T7p_3.3kbΔRBS, T7p_A-3C with pNL001; 50 μg/mL Kanamycin for dLacOperon) unless otherwise stated.

For the measurement of Venus-T7 RNAP and *lacZ* mRNA, the cells carrying the pNL001 plasmid were induced with appropriate anhydrotetracycline (aTc) concentration (0, 0.02, 0.05, 0.1, and 1 ng/mL) to control the expression level of Venus-T7 RNAP and 1mM isopropyl β-D-1-thiogalactopyranoside (IPTG) was added to achieve full expression of *lacZ* mRNA. The cells were grown at 37 ℃ for 4.5 h until reaching an $OD_{600}$ ~ 0.3.

##### mRNA Single-molecule FISH assay

*lacZ* mRNA measurements using smFISH were performed as previously reported (Skinner et al., 2013). The cell cultures were collected by centrifugation (4,500 g, 4 ℃, 5 min) and then were fixed in ice-cold 1× PBS with 3.7 % formaldehyde for 30 min at room temperature (RT). After fixation, the cells were washed twice in ice-cold 1× PBS, permeabilized in 70 % ethanol for 1 h at RT and washed again in wash buffer (2× SSC, 25 % formamide in DEPC-treated water). The cells were then incubated overnight at 30 ℃ with fluorescently labeled probes in hybridization buffer (2× SSC, 25 % formamide, 10 % dextran sulfate, 2 mM ribonucleoside-vanadyl complex, 0.2 mg/mL BSA and 1 mg/mL *E. coli* tRNA in DEPC-treated water). Then, 10 μL of the cells in hybridization reaction were washed twice with the wash buffer and incubated for 30 min ~ 1h at 30 ℃. For imaging, 1 μL of the cells were placed between a coverslip and 3 % low-melting-temperature agarose gel pad (Lonza, #50111) prepared with a 1× PBS.

##### *lacZ* mRNA fluorescent probes

The DNA oligonucleotides modified with a 3' amine group were ordered from Biosearch Technologies. The sequences of *lacZ* FISH probes are listed in Table S2. The probes were labeled with the fluorescent dye, ATTO 594 (ATTO-TEC GmbH).

##### Microscopy

An inverted microscope (Olympus, IX-71) with a 100x oil-immersed objective (Olympus) and an EMCCD (Andor iXon DU897) were used. The phase-contrast images and two fluorescence images of Venus and ATTO 594 were acquired from multiple fields of view. To image Venus, FF03-510/20 (Semrock) (excitation), HQ550/50m (Chroma) (emission), and FF520-Di01-25×36 (Semrock) (dichroic mirror) were used. Excitation was provided by a 514 nm Ar-ion laser (Melles Griot 43 Series Ion Laser). For imaging *lacZ* mRNA labeled with ATTO 594, FF01-572/28-25 (Semrock) (excitation), FF01-641/75-25 (Semrock) (emission) and FF593-Di02-25x36 (Semrock) (dichroic mirror) were used. Excitation was provided by a 580 nm fiber laser (VFL-P-Series, MPB Communications Inc.). Metamorph software (Molecular Devices) was used to control the automated measurements and maintain focus during data acquisition.

##### Image analysis

Home-built software (MATLAB) was used for the image analysis. Phase-contrast images were used to segment cell area and shape. Cells within a range of cell length ($< 1.3$ μm) were used for the further data analysis to take into account the cell cycle difference between cells. The total Venus-T7 RNAP intensities of individual cells were extracted and recorded. The average fluorescence intensity (concentration) was determined by normalizing the total intensity to the cell area. Autofluorescence was measured in cells without the plasmid expressing Venus-T7 RNAP. We used Spatz cells (Skinner et al., 2013) in the ATTO 594 fluorescence images to quantify the *lacZ* mRNA numbers in individual cells. The dLacOperon was used as the negative control for *lacZ* mRNA measurements.

##### Noise analysis using random selection of cells

The concept of the analysis is the same as that reported in our previous study (Yang et al., 2014). We generated gamma distributions based on the Venus-T7 RNAP intensities in individual cells to obtain pre-determined means (μ) and standard deviations (σ). To prepare the total cell collection, we combined all the cells (~ 20,000) obtained from measurements performed at different aTc concentrations (Fig. 2a, Supplementary Fig. 4). We then randomly selected 300 cells satisfying a given gamma distribution: $p(x) = \frac{1}{b^a\Gamma(a)} x^{a-1} e^{-x/b}$ with $a = \mu^2/\sigma^2$ and $b = \sigma^2/\mu$, from the total cell collection. We set the bin size for the Venus-T7 RNAP intensity to σ/2, calculated the number of cells required for each bin to satisfy the given gamma distribution and randomly selected the required number of the cells for each bin from the total cell collection. We then calculated the mRNA noise and the squared coefficient of variance using the selected cells. We repeated this process 1,000 times and obtained the mean and standard deviation of the noise.

**mRNA lifetime measurements using qRT-PCR**

The cells were grown in the same medium as in the smFISH experiments with 10 ng/mL aTc for 4 h to fully induce Venus-T7 RNAP expression. When the $OD_{600}$ of the culture reached 0.3, the cells were concentrated and re-suspended in a medium with 1 mM IPTG to induce *lacZ* expression for 30 min at 37 ℃.

50 μL of the cells were withdrawn into 150 μL of ice-cooled RNAlater solution (Ambion, AM7020) before shutting off *lacZ* expression for the time-zero sample. Then, the cells were pelleted by centrifugation (6,000 g, 1 min, RT) and re-suspended with a pre-warmed IPTG-free medium to shut off the *lacZ* expression under T7 promoter at 37 ℃. 50 μL of the cells were withdrawn into 150 μL of ice-cooled RNAlater solution every 30. After sampling, the cells were incubated for 20 min on ice and 400 μL of ice-cooled M9 medium was added to cells and gently mixed by pipetting. The cells were collected by centrifugation (10,000 rpm, 4 ℃, 2 min) and then the pellets were re-suspended in 100 μL of ice-cooled lysozyme solution (100 mg/mL lysozyme (Sigma, L4919), 10 mM Tris-HCl pH 8.0, 0.1 mM EDTA) and vigorously vortexed for 20 s. Subsequently, 0.5 μL of 10 % SDS solution (Sigma, L3771) was added to the cell. The cells were incubated at RT for 5 min. Total RNA was extracted using a PureLink RNA Mini Kit (Ambion) with DNase digestion (Invitrogen).

The extracted RNA was converted to cDNA using Superscript III Reverse Transcriptase (Invitrogen) and RNaseOut Recombinant Ribonuclease Inhibitor (Invitrogen). qRT-PCR was performed to determine the *lacZ* mRNA levels using SYBR Green premix (Enzynomics) and the StepOne Real-Time PCR system (Applied Biosystems). The sequences of the primers used are follows; 5'-tgttccgtcatagcgataac-3,' 5'-ttgcactacgcgtactgtga-3. ' The results were plotted to determine the lifetime of *lacZ* mRNA (Supplementary Fig. 3).

**Measurement of *in vivo* transcription kinetics**

All the experimental processes including the cell growth, the preparation of cells, the imaging acquisition, and the analysis of data were performed as previously described (Yang et al., 2019).

**Table S1 Bacterial strains used in this study.**

| Strain | Genotype | Plasmid | Reporter | Reference |
|---|---|---|---|---|
| T7p_3.3kb | O1-$P_{T7}$-O1-*lacZ*-tandem Tpi | pNL001<br>*aTc controllable expression of Venus-T7 RNAP* | Venus-T7 RNAP, *lacZ* mRNA | (Yang et al., 2019; Yang et al., 2014) |
| T7p_3.3kbΔRBS | O1-$P_{T7}$-O1-*lacZΔ RBS*-tandem Tpi | pNL001<br>*aTc controllable expression of Venus-T7 RNAP* | Venus-T7 RNAP, *lacZ* mRNA | (Yang et al., 2019; Yang et al., 2014) |
| | | pNL003<br>*rhamnose inducible expression of eYFP-T7 RNAP, in vivo imaging of T7 RNAP* | eYFP-T7 RNAP | (Yang et al., 2019) |
| T7p_A-3C | O1-$P_{T7(A\text{-}3C)}$-O1-*lacZ*-tandem Tpi | pNL001<br>*aTc controllable expression of Venus-T7 RNAP* | Venus-T7 RNAP, *lacZ* mRNA | This study |
| dLacOperon<br>*Negative control for lacZ mRNA* | Δ*lacIZYA* | - | - | (Yang et al., 2019) |

**Table S2 *lacZ* mRNA FISH probes.**

| Name | Sequence (5'-3') | Name | Sequence |
|---|---|---|---|
| lacZ1 | gtgaatccgtaatcatggtc | lacZ26 | tttgatggaccatttcggca |
| lacZ2 | attaagttgggtaacgccag | lacZ27 | aagactgttacccatcgcgt |
| lacZ3 | attcaggctgcgcaactgtt | lacZ28 | tgccagtatttagcgaaacc |
| lacZ4 | agtatcggcctcaggaagat | lacZ29 | aaacggggatactgacgaaa |
| lacZ5 | aaccgtgcatctgccagttt | lacZ30 | tcggcgtatcgccaaaatca |
| lacZ6 | aatgtgagcgagtaacaacc | lacZ31 | ttcatacagaactggcgatc |
| lacZ7 | gtagccagctttcatcaaca | lacZ32 | tggtgttttgcttccgtcag |
| lacZ8 | aataattcgcgtctggcctt | lacZ33 | acggaactggaaaaactgct |
| lacZ9 | agatgaaacgccgagttaac | lacZ34 | tattcgctggtcacttcgat |
| lacZ10 | tttctccggcgcgtaaaaat | lacZ35 | gttatcgctatgacggaaca |
| lacZ11 | atcttccagataactgccgt | lacZ36 | gttcaggcagttcaatcaac |
| lacZ12 | aacgagacgtcacggaaaat | lacZ37 | ttgcactacgcgtactgtga |
| lacZ13 | gctgatttgtgtagtcggtt | lacZ38 | agcgtcacactgaggttttc |
| lacZ14 | aactgttacccgtaggtagt | lacZ39 | cggttaaattgccaacgctt |
| lacZ15 | atagagattcgggatttcgg | lacZ40 | ctgtgaaagaaagcctgact |
| lacZ16 | ttctgcttcaatcagcgtgc | lacZ41 | tacgccaatgtcgttatcca |
| lacZ17 | accattttcaatccgcacct | lacZ42 | taaggttttcccctgatgct |
| lacZ18 | ttaacgcctcgaatcagcaa | lacZ43 | atcaatccggtaggttttcc |
| lacZ19 | tctgctcatccatgacctga | lacZ44 | gtaatcgccatttgaccact |
| lacZ20 | cacggcgttaaagttgttct | lacZ45 | agttttcttgcggccctaat |
| lacZ21 | ttcatccaccacatacaggc | lacZ46 | atgtctgacaatggcagatc |
| lacZ22 | tgccgtgggtttcaatattg | lacZ47 | ataattcaattcgcgcgtcc |
| lacZ23 | atacagcgcgtcgtgattag | lacZ48 | tgatgttgaactggaagtcg |
| lacZ24 | gatcgacagatttgatccag | lacZ49 | attcagccatgtgccttctt |
| lacZ25 | aaataatatcggtggccgtg | lacZ50 | aatccccatatggaaaccgt |

#### Supplementary Figures

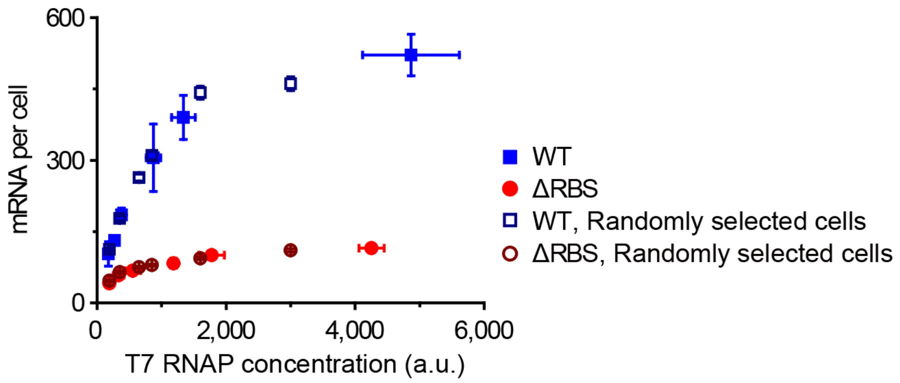


**Supplementary Figure 1.** To confirm whether cells with similar T7 RNAP concentrations express similar mRNA levels, regardless of the aTc induction conditions, we divided the total cell distribution into several subsets depending on the T7 RNAP concentration. The mean mRNA copy numbers are plotted against the T7 RNAP concentration in each subset (open symbols). The filled red circles and blue squares represent the mean *lacZ* mRNA expression levels of WT and ΔRBS, respectively, at each aTc concentration shown in Fig. 3b.

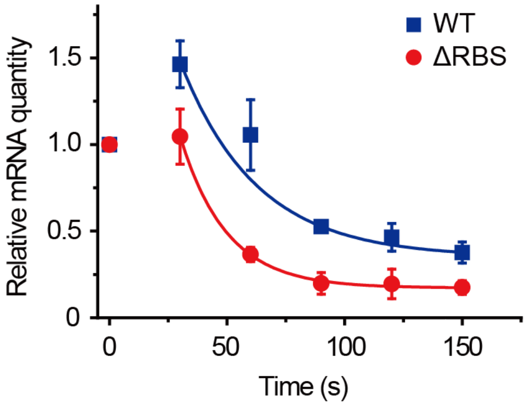


**Supplementary Figure 2. mRNA lifetime measurements.**

The lifetime of the *lacZ* mRNA with and without the RBS. Upon turning off transcription driven by the T7 promoter, the relative *lacZ* mRNA level was measured by qRT-PCR, at each time point. The lines are fits to $y=A+B \exp(-x/t)$, and $t$ is mRNA lifetime. The lifetime of the WT was 56 ± 6 s and of the ΔRBS was 18 ± 4 s. Error bars were s.d. from two independent experiments.

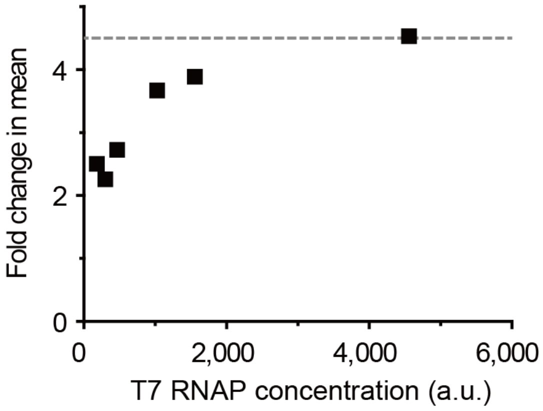


**Supplementary Figure 3. Fold change in mean mRNA level.**

The fold change in mean is obtained from the mean mRNA level of WT divided by that of ΔRBS at each aTc concentration and then is plotted as a function of T7 RNAP concentration. At the saturation level of T7 RNAP concentration, the fold change is 4.5, indicated as the dotted line.

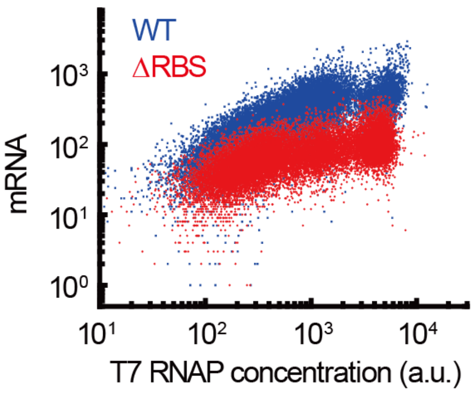


**Supplementary Figure 4. RNAP concentration dependence of mRNA level.**

Each spot represents a single cell with its T7 RNAP concentration and mRNA levels on the x and y axes, respectively. Total cells are combined from the cells obtained at the each aTc concentration condition. Blue spots representing WT reproduced Fig. 2a for comparison.

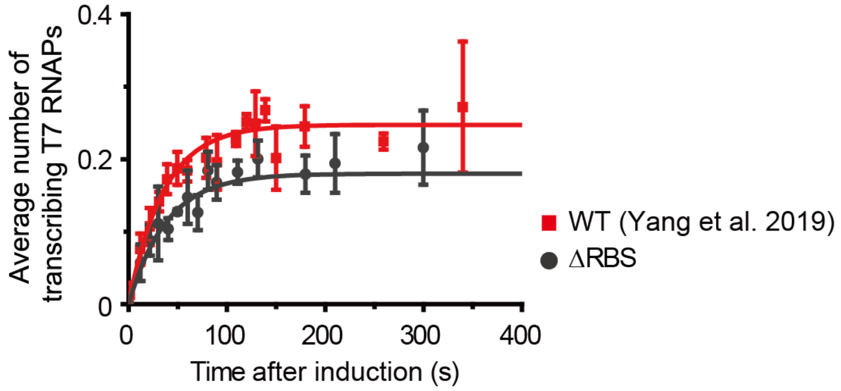


**Supplementary Figure 5. Ribosome binding onto mRNA increases transcription on-rate.** T7 RNAP transcriptional kinetics with (red squares, adapted from Yang *et al*. 2019 Nat Comm for comparison) and without (gray circles) RBS. The transcription elongation rates are 84 ± 8 bp $s^{-1}$ with RBS and 83 ± 6 bp $s^{-1}$ without RBS. The transcription on times are 131 ± 10 s with RBS and 197 ± 24 s without RBS (the ratio is ~ 0.66) when the average number of T7 RNAP in a cell was ~ 35. All error bars were obtained from three independent experiments.
